# Inferring in vivo murine cerebrospinal fluid flow using artificial intelligence velocimetry with moving boundaries and uncertainty quantification

**DOI:** 10.1101/2024.08.29.610340

**Authors:** Juan Diego Toscano, Chenxi Wu, Antonio Ladrón-de-Guevara, Ting Du, Maiken Nedergaard, Douglas H. Kelley, George Em Karniadakis, Kimberly A. S. Boster

## Abstract

Cerebrospinal fluid (CSF) flow is crucial for clearing metabolic waste from the brain, a process whose dysregulation is linked to neurodegenerative diseases like Alzheimer’s. Traditional approaches like particle tracking velocimetry (PTV) are limited by their reliance on single-plane two-dimensional measurements, which fail to capture the complex dynamics of CSF flow fully. To overcome these limitations, we employ Artificial Intelligence Velocimetry (AIV) to reconstruct three-dimensional velocities, infer pressure and wall shear stress, and quantify flow rates. Given the experimental nature of the data and inherent variability in biological systems, robust uncertainty quantification (UQ) is essential. Towards this end, we have modified the baseline AIV architecture to address aleatoric uncertainty caused by noisy experimental data, enhancing our measurement refinement capabilities. We also implement UQ for the model and epistemic uncertainties arising from the governing equations and network representation. Toward this end, we test multiple governing laws, representation models, and initializations. Our approach not only advances the accuracy of CSF flow quantification but also can be adapted to other applications that use physics-informed machine learning to reconstruct fields from experimental data, providing a versatile tool for inverse problems.

## 1. Introduction

The cerebrospinal fluid (CSF) flow in the brain plays an important role in transporting solutes, including clearing metabolic waste whose buildup has been linked to Alzheimer’s disease [1, 2]. This system of solute transport is often referred to as the glymphatic system and has been shown to be altered with aging in mice and humans [3, 4]. It is hypothesized to be changed with functional hyperemia [5], under different anesthetic states [6], and in pathological scenarios such as stroke [7], hypertension [8], cerebral amyloid angiopathy [9], and cerebral small vessel disease [10]. CSF is delivered rapidly around the brain’s surface in a network of channels adjacent to surface (pial) arteries called perivascular spaces (PVSs) before it enters deep into the brain tissue, where solute exchange occurs. Quantifying the flow of CSF is essential to understanding glymphatic function, including building models of glymphatic flow, designing drug delivery systems, and, eventually, developing interventions to treat glymphatic-related pathologies. This work focuses on quantifying the flow of CSF in surface PVSs.

Traditionally, CSF flow has been quantified using particle tracking velocimetry (PTV) in surface PVSs, which only provides velocity data within a single plane and can lead to misrepresenting the volumetric flow rate, which is a more pertinent metric than velocity. To address these challenges, physics-informed machine learning (PIML) has been explored as an alternative for modeling complex biological flows [11–13]. The PIML approach uses a representation model, such as multilayer perceptrons or Kolmogorov-Arnold networks, to approximate the solution of ordinary/partial differential equations (ODE/PDE) by minimizing a multi-objective loss function that attempts to fit any observable data while satisfying the underlying physical laws [11]. PIML flexibility makes it suitable for solving forward and inverse problems[12, 14], and it has been shown in [15] that, for some linear problems such as Stokes flow, any neural network that sufficiently minimizes the loss function is a good approximation to the true solution. Despite their advantages, PIML effectiveness can be compromised by the nonlinear and chaotic nature of biological systems, so enhanced neural network architectures, transformational methods, and adaptive strategies are required to maintain accuracy and reliability [16–26]. For instance, Boster et al. [13] used AIV, enhanced with learning rate annealing[21], to model the CSF flow from sparse 2D data and boundary conditions, obtaining volumetric flow rates and other dynamical parameters. However, one limitation of their study was the assumption of stationary boundaries, given the dynamic nature of boundary movements driving CSF flow [8, 27, 28]. Despite the impressive reconstruction and inference capabilities of AIV [16, 22, 29], neural networks are poor at quantifying predictive uncertainty and tend to produce overconfident predictions [30], which can be harmful to biological or medical applications. Additionally, complex deep-learning models usually lack interpretability. Thus, the integration of uncertainty quantification (UQ) with deep learning (DL) has emerged as the potential solution to improve the safety, interpretability, and reliability of neural network predictions [31]. Psaros et al. [32] identified that the uncertainties in PIML are sourced in several factors, which can be roughly categorized into aleatoric, stemming from noisy or sparse data, epistemic, arising from the representation model (i.e., architecture, initialization, etc.), and model uncertainty stemming from the governing equations.

To address the highlighted issues effectively, we extend the AIV framework proposed in [13] by incorporating moving boundary conditions and quantifying the uncertainties in quantities of interest, such as flow rate, pressure gradient, and wall shear stress. To systematically quantify aleatoric uncertainty, we introduce AIV-NLL, a refined scientific machine-learning approach that combines AIV and a negative-log likelihood (NLL) criterion [30, 33, 34] to infer flow fields and their uncertainties associated with noisy experimental data. AIV-NLL enhances the model interpretability and provides information on the confidence of the AIV framework in making predictions. To quantify model and epistemic uncertainties, we employ an ensemble of models (EoM) [30]. In this approach, multiple models are independently trained to perform the same task. The predictive probabilities from these models are then approximated by a Gaussian distribution, where the ensemble’s mean and variance are used to represent the mean and variance of this distribution, respectively, thus providing uncertainty estimates [30]. This method offers a robust and scalable framework for understanding how variations in the model representation — such as parameter initialization, model architecture, and governing equations — impact the results. It is important to note that when the EoM is applied to variations in parameter initialization, it aligns with the deep ensemble method [30].

Through these strategies, our work aims to enhance the accuracy and reliability of CSF flow quantification in surface PVSs and deepen our understanding of our model capabilities by explicitly addressing and quantifying the uncertainties associated with each component of the modeling process. We show that the flow fields inferred by AIV, and precisely the volume flow rate, pressure gradient, and shear stress at the wall, are relatively robust to variations to the governing equations modeling the flow, significant changes to the network architecture, and the initial conditions; in short, the combined aleatoric, epistemic and model uncertainties are less than 30%.

## 2. Problem Description

To obtain the experimental data, Boster et al. [13] injected one-micron fluorescent microspheres and dyes into the CSF, enabling visualization and quantification of the flow velocity within the PVS. The position and velocity of the microsphere particles over time were quantified using particle tracking velocimetry, and the 3D PVS geometry was reconstructed from a 3D volume scan, as illustrated in Figure 1(A). In this study, we also add PVS domain boundary motion, which we infer from the contraction of the adjacent vessel. To enhance the resolution of the PTV data, we consolidated PTV data from multiple cardiac cycles into a single (phase-merged) cardiac cycle of *T* = 0.303*s*. Finally, to evaluate the PDE residuals required to enforce the governing equation during training, we generated collocation points by shrinking and combining the boundary conditions, as shown in Figure A.9. This strategy allows us to control the spatial distribution of points within our domain.

**Figure 1.**
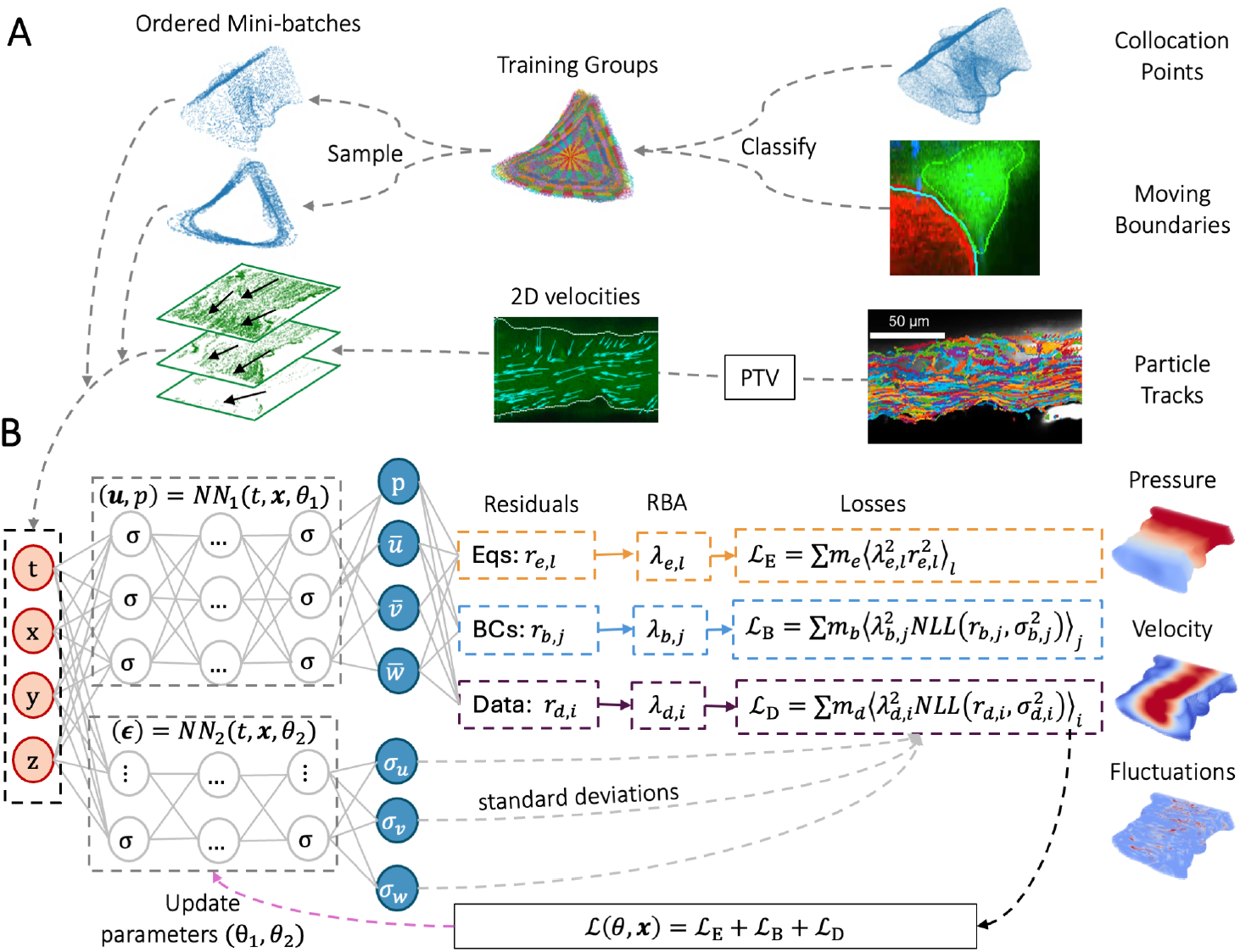
Schematic diagram of AIV-NLL. (A) The model is trained using in vivo particle tracks, moving boundary conditions (MBCs), and collocation points (i.e., points used to evaluate the PDE) sampled in the perivascular space (PVS). The PVS domain is represented in green, and the adjacent artery is represented in red. MBCs and collocation points are clustered into training groups, from which ordered mini-batches are obtained. 2D velocity measurements are obtained by particle tracking velocimetry (PTV). (B) At each iteration, the point coordinates are fed into two neural networks. The outputs from the first neural network are the pressure and mean velocity fields, while the second network predicts the standard deviation of each velocity component. Residuals for PTV, MBCs, and equations are calculated. Residual-based attention (RBA) multipliers are updated using an exponentially weighted moving average of the residuals. The scaled residuals form each loss subterm, namely the negative log-likelihood criterion for PTV and MBCs and the mean-squared error for the equation residuals. The total loss updates the parameters, resulting in continuous and differentiable pressure, mean velocity, and standard deviation fields.

Based on these experimental data, we follow [13, 16] and use AIV to infer three-dimensional (3D) velocities and pressure from two-dimensional (2D) particle tracking velocimetry (PTV) data and 3D moving boundary conditions. We train our model by optimizing a multi-objective loss function (L) that minimizes the point-wise error (i.e., residuals) of the velocity data (*r*_*d*_), boundary conditions (*r*_*b*_), and governing equations (*r*_*e*_) (see Figure 1(B)). To deal with the local imbalances related to optimizing ℒ, we use residual-based attention (RBA) weights (*λ*_*i*_) as local multipliers to balance the point-wise errors, enabling a uniform convergence along the analyzed domain [17, 35]. Using this approach, we obtain continuous and differentiable flow fields that we use to compute quantities of interest such as flow rate, pressure gradient, and wall shear stress, which are relevant to understanding the fluid dynamics involved in glymphatic flows. Despite the impressive reconstruction and inference capabilities of AIV [16, 22, 29], neural networks (NNs) often struggle with quantifying prediction uncertainty. To address these challenges, we quantify uncertainty in three quantities of interest: flow rate, pressure gradient, and wall shear stress. To capture aleatoric uncertainty, we train a second neural network that predicts the standard deviations of each velocity component, learning these values using a negative log-likelihood (NLL) criterion [34] (see Figure 1(B)). For epistemic and model uncertainties, we employ an ensemble of models [30], where multiple models are trained to perform the same task. Specifically, to assess model uncertainty, we train six networks constrained by different systems (e.g., PDEs, BCs, or additional assumptions) that ideally govern the CSF flow and compute the uncertainty in the three quantities of interest. Following a similar approach, we identify epistemic uncertainty due to parameter initialization and neural network enhancements.

## 3. Methods

### 3.1. Experimental data collection

#### 3.1.1. Mouse surgery, imaging, and particle tracking velocimetry

A cranial window was installed, and one-micron fluorescent particles and dye were injected into the CSF in the cisterna magna (Fig. 2a) of a mouse in vivo. A dextran dye was injected intravenously to indicate the location of the blood vessel. The PVS geometry was reconstructed from a 3D volume scan (see Fig. 1a). The motion of the fluorescent particles in a single plane (plane A) was tracked using particle tracking velocimetry (PTV) (Fig. 2b, c). Vessel motion was obtained from time-series images at a different cortical depth (plane B) where more of the vessel was visible. We use Cartesian coordinates (*x,y,z*), where *z* aligns with the depth direction on the microscope and *y* roughly aligns with the mean flow direction. Further details regarding mouse surgery, anesthesia, imaging, and particle tracking are described by Boster et al. [13].

**Figure 2.**
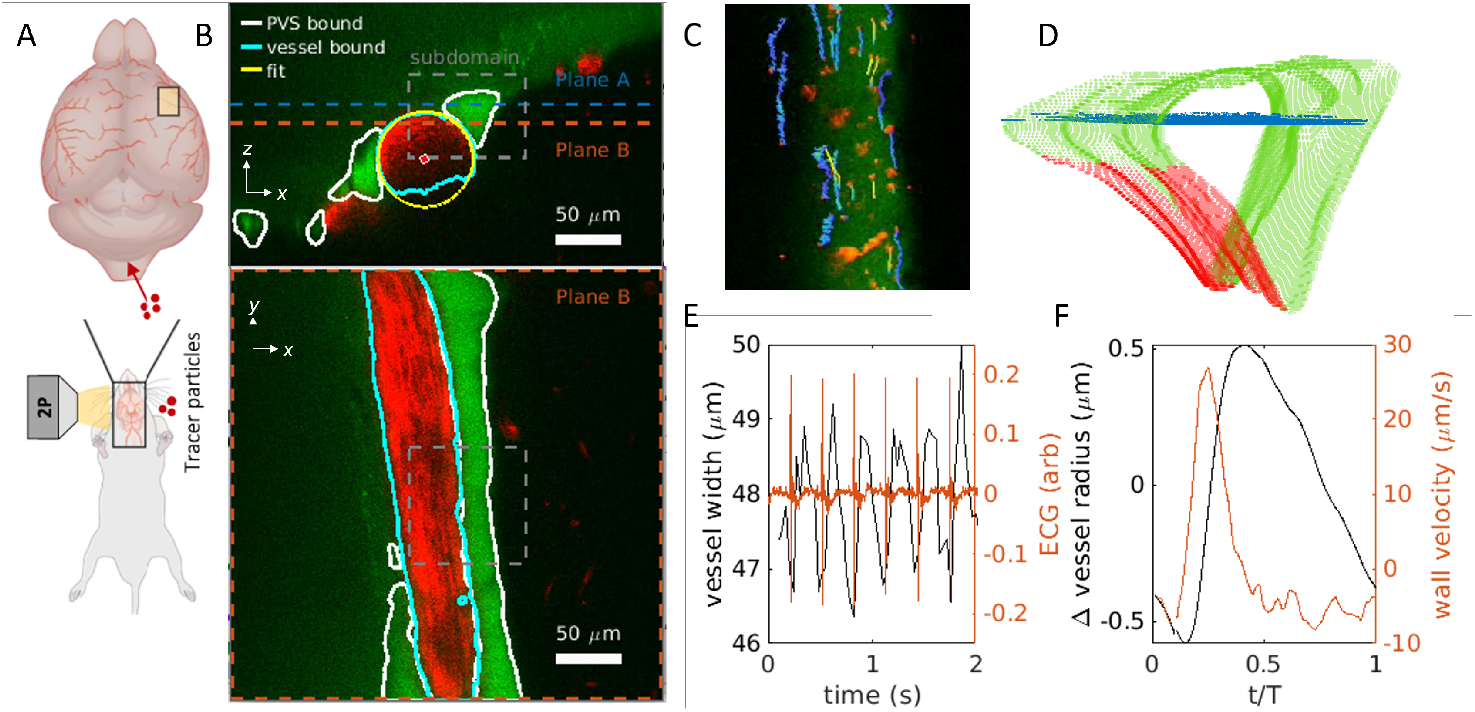
Overview of training data. (a) Fluorescent particles and dyes are injected into the mouse and imaged with two-photon microscopy. (b) A two-photon microscopy image from a volumetric scan shows an axial view of the vessel (red) and surrounding PVS (green). High temporal resolution imaging was acquired at planes A and B. (c) Example particle tracks and (d) 3D boundaries of the PVS, with the moving portion of the boundary indicated in red and the particles acquired from plane A. (e) Vessel diameter was measured and then averaged over the cardiac cycle, as determined from simultaneous electrocardiogram measurements. The vessel radius over a cardiac cycle was then inferred by assuming axisymmetric vessel pulsations.

#### 3.1.2. Moving Boundaries

Arterial vessels pulse in synchrony with the cardiac cycle, and since the vessel wall is directly adjacent to one side of the PVS, the vessel pulsatility moves the PVS boundary adjacent to the vessel. Other regions of the PVS may also pulse with the cardiac cycle. Still, that motion is not well characterized and cannot be determined from the time-series images of the PVS because the PVS boundaries are not very distinct since they are only indicated by the presence of a dye in the PVS lumen. Additionally, even if the boundaries were clearly indicated (as possible with a transgenic mouse labeled with a fluorescent protein that indicated the PVS boundaries), that would still only provide motion in a single plane, and it is unclear how to infer motion in three dimensions. In contrast, the vessel wall boundaries are more distinct, and we can infer motion in three dimensions by assuming axisymmetric vessel pulsatility. Therefore, we infer the motion of the PVS boundaries adjacent to the vessel based on the vessel wall motion and assume stationary boundaries elsewhere. The following paragraph describes the procedure used to determine this motion.

Time-series images of the blood vessel and PVS was obtained from a plane intersecting the vessel (plane B in Fig. 2b). The vessel width in plane B was obtained using a Canny edge detection algorithm (the results shown in Fig. 2e). The time-series width measurement was phase-averaged over the cardiac cycle based on simultaneous ECG recordings. Based on the distance of plane B from the vessel centerline, the phase-averaged vessel width was converted into a vessel radius, and the vessel wall velocity was calculated by taking the derivative of the radius with respect to time (Fig. 2f). The three-dimensional boundaries of the vessel were determined from two-photon microscopy. We fit a circle to the location of the vessel boundaries in axial slices spaced 0.6480 µm apart, as described by Raicevic et al. [36] (Fig. 2b). The contrast along the bottom edge of the vessel is much worse than the rest of the vessel because red blood cells attenuate the signal intensity, so we excluded points on the bottom edge of the vessel. From the fit circle, we determined the vessel centerline and calculated the position and velocity of the vessel boundary points, assuming that the vessel pulsation was axisymmetric. In order to decide which points on the PVS boundary were adjacent to the vessel, we dilated the vessel by 4 µm. We classified all boundary points inside the dilated vessel as moving and all other boundary points as stationary (Fig 2d). The moving boundary points were assigned the same velocity magnitude as the vessel, and the velocity direction was calculated with respect to the vessel’s centerline.

### 3.2. Underlying Physical Laws

The flow of CSF in the PVS obeys the Navier-Stokes (NS) equations. Therefore, we follow [12, 37, 38] and define momentum and mass conservation equations as:

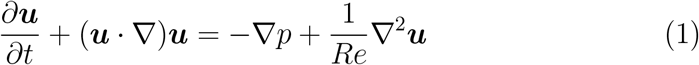

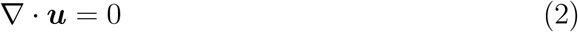

where ***u*** = (*u, v, w*) is the nondimensional velocity field, *p* is the nondimensional pressure, and *Re* = 1.1 × 10^−3^ is the Reynolds number [13].

Since *Re* ≪1, the inertial terms (i.e., the terms 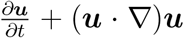 on the left-hand side of equation (1)) are negligible. This theoretical assumption was verified in [29]. Consequently, in this study, we model this system using Stokes flow:

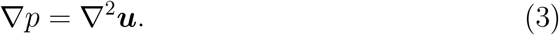

This simplification eliminates the nonlinearities of the NS equation, which enables the use of the negative log-likelihood criterion [30, 33, 34].

#### 3.2.1. Quantities of interest

##### Pressure gradient

The pressure gradient within the PVS is exciting from a fluid dynamics standpoint because it drives the glymphatic flow. Once our model is trained, we can calculate the axial pressure gradient 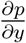 using automatic differentiation.

##### Volume flow rate

The flow of CSF is typically the dominant mass transport mechanism in surface PVSs and has most often been quantified using the mean velocity (measured by PTV). However, the mean velocity calculated from PTV will not reflect the average PVS velocity if the PTV imaging plane is not close to the center of the PVS or if the measurement locations are not uniformly distributed in the PVS, as shown by Boster et al. [13]. A better metric is the volume flow rate *Q*, which we calculate from the velocity component *v* by sampling on a square grid (resolution 0.648 µm):

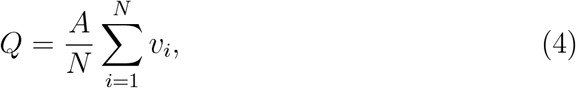

where *A* is the cross-sectional area and *i* indexes the *N* individual velocity inferences.

##### Shear stress

The shear stress is defined as

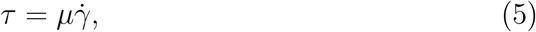

where 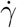 is the shear rate, calculated using the strain rate tensor **E**:

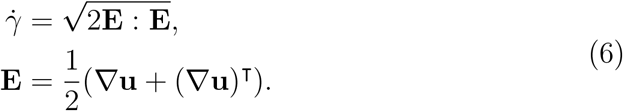

As defined, *τ* is a scalar magnitude accounting for all shear components, not just the component-oriented downstream along the wall (often referred to as “wall shear stress”). We expect that the magnitude, rather than a single component, is likely the relevant quantity for aggregation and mechanical signaling. While the role of PVS wall shear stress in glymphatic flows is not known, wall shear stress plays a significant role in cardiovascular flows, and quantifying the shear stress is the first step in elucidating its importance.

### 3.3. Uncertainty Quantification

Given a set of paired noisy observations 𝒟 = {***x***_*i*_, ***u***_*i*_}, our goal is to construct the probability distribution *p*(***u***|***x***, 𝒟, ℋ) for predicting ***u*** at any new location ***x***, where ***x***_*i*_ = (*t*_*i*_, *x*_*i*_, *y*_*i*_, *z*_*i*_) denotes the time and three-dimensional coordinates and denotes a set of assumptions such as neural network architecture, governing equations, choice of the optimizer, etc. Following [32], we assume that ***u***(***x***_*i*_) is generated by a data-generating process comprising a deterministic component 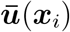 and an additive noise ***u***^*′*^(***x***_*i*_), representing aleatoric uncertainty. For modeling this process, we assume a likelihood function *p*(***u***|***x***, *θ*), where *θ* are the parameters to be inferred from the data, introducing epistemic uncertainty regarding the representation model, and model uncertainty stemming from the governing equations [32, 39–41].

We employ two approaches to quantify the uncertainty related to our problem. To measure the aleatoric uncertainty, we combine AIV with the negative log-likelihood method [33]. On the other hand, we use the ensemble of models (EoM) to compute epistemic and model uncertainties [30] due to the parameter initialization, model enhancements, and governing equations. We chose EoM over traditional Bayesian methods due to the computational challenges and potential inaccuracies of Bayesian neural networks (BNNs). While BNNs use a prior distribution on parameters to compute a posterior for predictive uncertainty, the need for approximations often compromises the quality of uncertainty quantification [42, 43]. These approximations can yield unreliable predictive uncertainties [44]. Furthermore, traditional Bayesian methods, including Bayesian Model Averaging (BMA), are typically slower to train and more challenging to implement than ensemble methods. Ensembles provide a robust mechanism for capturing a broader spectrum of model behaviors, particularly advantageous when the actual model may not align with the predefined hypothesis class [30]. Our study leverages EoM to explore their effectiveness in estimating predictive uncertainty, demonstrating their practical advantages in modeling complex systems.

#### 3.3.1 Aleatoric Uncertainty

We use NLL to quantify the aleatoric uncertainty. Towards this end, we assume that the velocities ***u*** = (*u, v, w*) can be decomposed into mean fields 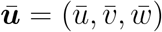 and fluctuations from the mean ***u***^*′*^ = (*u*^*′*^, *v*^*′*^, *w*^*′*^), namely:

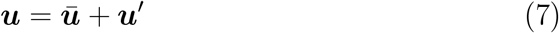

In this study, we assume that the mean fields follow the Stokes (S) equations and that the fluctuations are sampled from a normal distribution ***u***^*′*^ ∼ 𝒩(0, ***σ***) centered on zero and with standard deviation ***σ*** = (*σ*_*u*_, *σ*_*v*_, *σ*_*w*_). Under these assumptions, we can learn the data-driven mean velocities 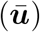 and their standard deviations (***σ***) using the NLL criterion proposed in [30, 33, 34].The NLL is described as follows:

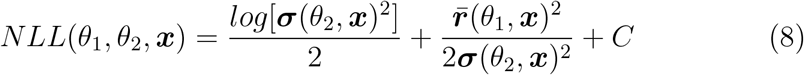

where ***x*** = (*t, x, y, z*) are the non-dimensional inputs (i.e., time and position), and *θ*_1_, and *θ*_2_ are two sets of trainable parameters from two independent neural networks (see Figure 1(B)). The residuals 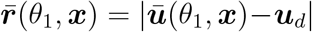 are the absolute difference between the data ***u***_*d*_ and the predicted mean field 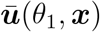. Finally, *C* is a positive constant and ***σ***(*θ*_2_, ***x***) = (*σ*_*u*_(*θ*_2_, ***x***), *σ*_*v*_(*θ*_2_, ***x***), *σ*_*w*_(*θ*_2_, ***x***)) are the predicted standard deviations. Notice that the residual from the data is scaled by the corresponding standard deviation; therefore, it behaves as an attention mask that helps the model fit the data better. Thus, this technique can also be considered as an optimization method.

We learn the mean fields *u, v, w* and their corresponding standard deviations *σ*_*u*_, *σ*_*v*_, *σ*_*w*_ by optimizing a combined loss function that minimizes errors from data, boundary conditions, and equations. The data loss (ℒ_*D*_) controls the mismatch between the network prediction and experimental observations and is explicitly defined as:

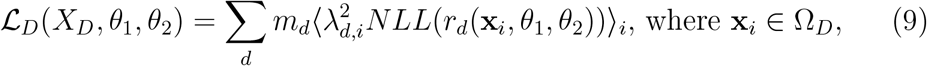

where ⟨·⟩_*i*_ is the mean operator and **x**_*i*_ = (*t*_*i*_, *x*_*i*_, *y*_*i*_, *z*_*i*_) is the *i*^th^ point from batch *X*_*D*_ which was selected from the data domain Ω_*D*_. The index *d* = {*u, v, w*} identifies the specific variables constrained in the loss, where *u, v, w* represent the velocities in the *x, y, z* directions, respectively. The residual (i.e., point-wise error) 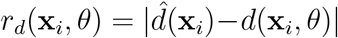 quantifies the difference between the experimental observation 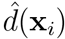 and the network prediction *d*(**x**_*i*_, *θ*) at point **x**_*i*_ ∈ Ω_*D*_. We use residual-based attention weights (RBA) [16] as local multipliers (*λ*_*d,i*_) to balance the point-wise contribution of the residual *r*_*d*_(**x**_*i*_, *θ*) and global weights (*m*_*d*_) to scale the averaged value of component *d*.

Similarly, the boundary conditions are imposed by the boundary loss ℒ_*B*_:

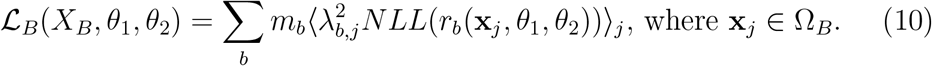

Here, Ω_*B*_ is the boundary domain corresponding to the walls of the PVS channel. To impose the no-slip boundary conditions, we set *b* = {*u, v, w*}, with residuals *r*_*b*_(*x*_*j*_, *θ*) = |*b*(*x*_*j*_, *θ*)|, RBA weights *λ*_*b,j*_, and global weights *m*_*b*_. To enforce our governing equations, we rewrite equations 1 and 2 in their residual form and impose them iteratively by minimizing the loss function described in equation 11.

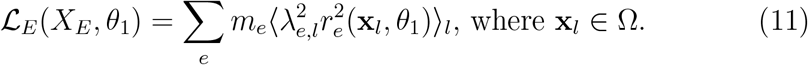

Here, *e* = {*M*_*x*_, *M*_*y*_, *M*_*z*_, *DF*} identifies the residuals *M*_*x,y,z*_ from the momentum equations in the {*x, y, z*}directions, respectively (i.e., equation 3), and the residual *DF* from conservation of mass (i.e., equation 2). We define the residual for subcomponent *e* and point *x*_*l*_ ∈ Ω as *r*_*e*_(*x*_*l*_, *θ*) = |*e*(*x*_*l*_, *θ*)|, and balance its point-wise and averaged contribution to ℒ_*E*_ using RBA as local multipliers *λ*_*e,l*_ and global weights *m*_*e*_, respectively.

To stabilize the training process [16], ordered batches are generated based on the spatial and temporal information (See Figure 1(A)). Finally, to speed up convergence, we re-parameterize our network using weight normalization (WN) [45]. The remaining details about the implementation are shown in Appendix A.

##### Volume flow rate aleatoric uncertainty

Once the model is trained, we obtain continuous and differentiable velocity fields. Then, following [13], we compute the volume flow rate at planes near the inlet and outlet using equation 4. Quantifying the uncertainties of this derived quantity is not straightforward since it is unclear how the measurements are related. Nevertheless, by assuming that the values are perfectly correlated, we can compute an upper bound:

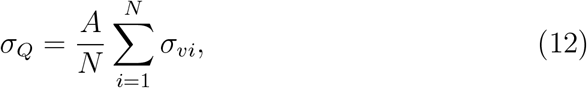

where *A* is the cross-sectional area, *N* is the number of points in the grid, and *σ*_*vi*_ is the predicted standard deviation related to the *v* velocity for the *i*^th^ point.

#### 3.3.2. Epistemic Uncertainty

We quantify the epistemic uncertainty using EoM [30]. This method leverages the variability among different models trained on the same data, making it robust as it captures how different model configurations and training paths can lead to different predictions, thereby reflecting the uncertainty. Towards this end, we assume that differences in the final prediction due to the representation model can be sourced from two main factors: parameter initialization and model enhancements, as better optimization methods can lead to different outcomes and introduce uncertainty.

To quantify the uncertainty from these three components, we train *M* independent neural networks, varying one source of uncertainty at a time. We predict the corresponding flow fields and compute the quantities of interest. For ease of predicting predictive probabilities, we follow [30] and approximate the ensemble prediction as a Gaussian distribution whose mean and variance are, respectively, the mean and variance of the mixture:

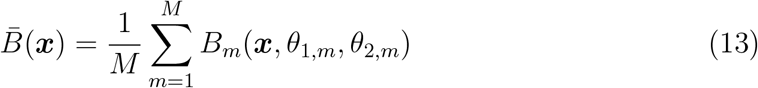

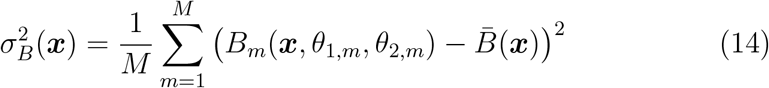

where 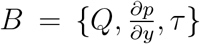 are the quantities of interest, namely, flow rate *Q*, pressure gradient 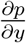 and wall shear stress *τ*.

##### Initialization

Following [30], we independently train *M* = 5 identical models initializing them with five different seeds. The initialization plays a significant role since it can lead to different local minima. For this case, we use Stokes flow with moving boundary conditions, weight normalization, RBA, and negative log-likelihood.

##### Model enhancements

To analyze the impact of the model enhancements, we initialize our neural networks identically and train them using Stokes flow (equations 3 and 2) with moving boundary conditions. Then we consider the model enhancements, namely weight normalization (WN), residual-based attention weights (RBA), and negative log-likelihood (NLL), and analyze the following combinations:

1. WN+RBA.
2. WN+NLL.
3. RBA+NLL.
4. WN+RBA+NLL.
5. NLL

#### 3.3.3. Model Uncertainty

To assess the impact of the physical model (i.e., model uncertainty), we initialize *M* = 6 models using the same random seed, weight normalization, and RBA. Each of these models is trained independently using one of the following physical systems (i.e., governing equations, boundary conditions, and assumptions):

1. NS equations (i.e., equations 1 and 2) with moving boundary conditions and NLL (NS+MBCs+NLL). Recall that under NLL, the velocities are decomposed into mean and fluctuations, with only the mean fields following the governing equations.
2. NS equations with moving boundary conditions (NS+MBCs).
3. NS equations with fixed boundary conditions (NS).
4. S flow (i.e., equations 3 and 2) with moving boundary conditions and NLL (S+MBCs+NLL).
5. S flow with moving boundary conditions (S+MBCs).
6. S flow with fixed boundary conditions (S).

Notice that the same PDE with different boundary conditions induces different solutions and can thus be considered a different model. Similarly, when using NLL, we decompose the velocity into mean and fluctuations and assume that our mean fields satisfy the governing equation. Therefore, the same PDE under different assumptions can be considered a different physical model.

#### 3.3.4. Combined uncertainty quantification

We calculate the combined uncertainty as follows:

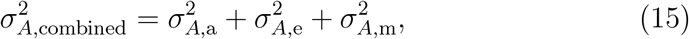

where 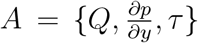, are the quantities of interest, *σ*_*A,α*_ is the standard deviation calculated using equation 14, *α* is a subscript *α* = {a, e, m} used to identify the aleatoric, epistemic and model uncertainty, respectively. The epistemic uncertainty is defined as follows:

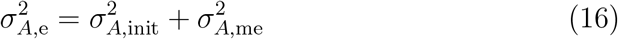

here the subscript *e* = {init, me} identifies the uncertainty from the parameter initialization and model enhancement, respectively.

## 4. Results

### 4.1. Aleatoric Uncertainty

As shown in Figure 3(A), we can obtain high-resolution 3D pressure, mean velocities, and velocity standard deviations. Notice that, using this approach, we can get the standard deviation at any point inside the domain, enabling us to compute the aleatoric uncertainty modeled as heteroscedastic noise [30]. Following [13], we assess the capability of our method to reconstruct the velocity fields, and we compare the model predictions on the validation dataset (i.e., 70% of the measured velocity data). Figure 3(B) shows that the distribution of the PTV velocities agrees well with the values inferred by the neural network. Similarly, Figure 3(C) shows the model predictions of the three velocity components on the boundaries. Figure 3(D) shows that the predicted uncertainty is higher in the interface between the moving and static regions. Finally, Figure 3(E) shows that the distributions of the data and the predictions compare well. The relative *L*^2^ errors (equation A.8) for the moving boundary conditions for *u* and *w* are 10.39% and 10.01%. Finally, the relative *L*^2^ errors for the PTV data on the validation dataset for *u* and *v* are 23.95% and 18.63%, respectively.

**Figure 3.**
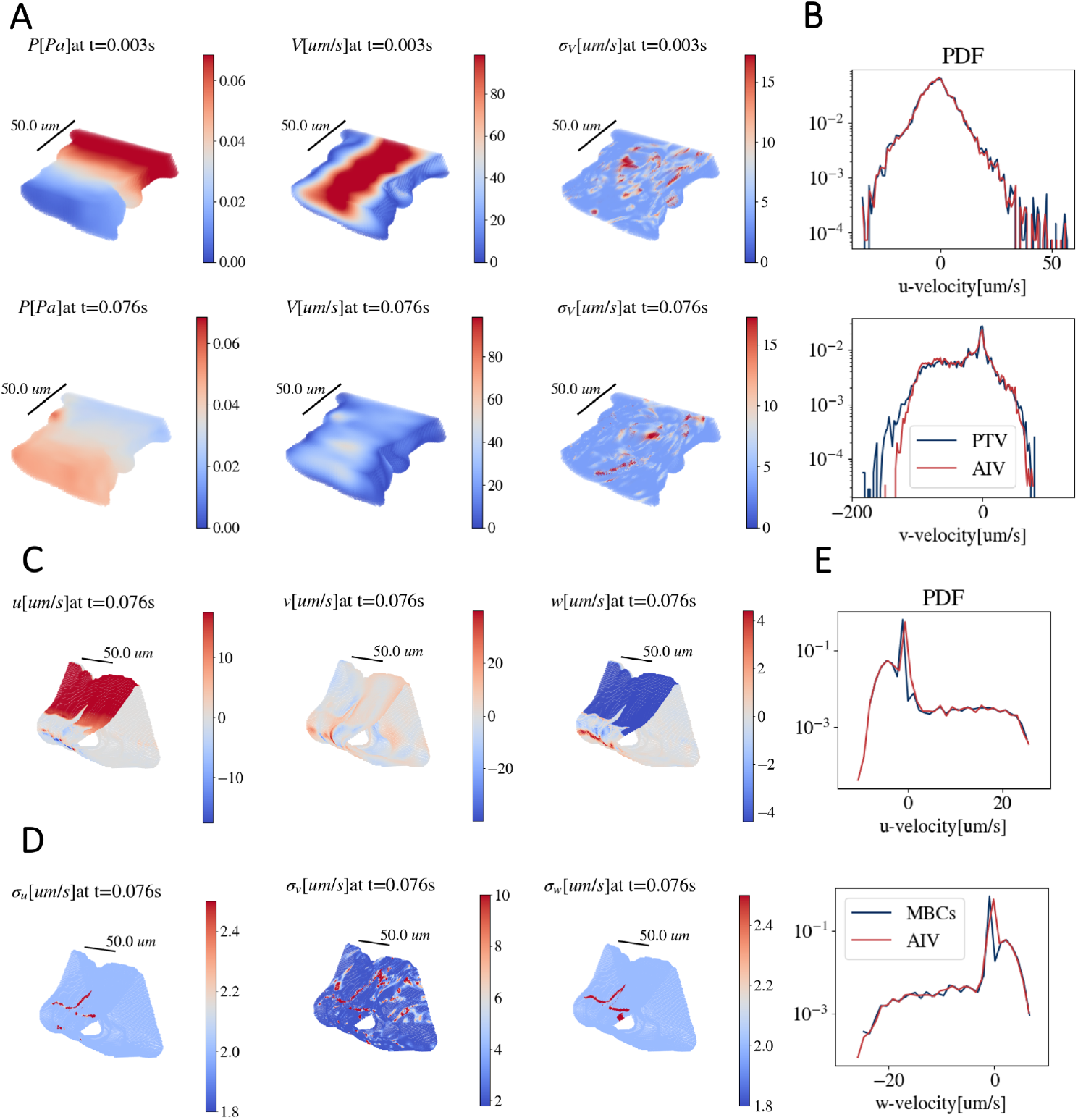
(A) Predicted pressure *P*, mean velocity magnitude *V*, and standard deviation magnitude *σ*_*V*_ on the lower half domain, i.e., *z* ≤ 0.5(*z*_*min*_ + *z*_*max*_) at two representative times (first and second row). (B) Probability density functions (PDFs) of two velocity components from PTV data (blue) and predicted mean velocities at particle locations (red) in the validation dataset. Predicted mean velocity fields (C) and their corresponding standard deviations (D) on the boundaries at a representative time. (E) Probability density functions of two velocity components, from MBCs data (blue) and predicted velocities (red).

Since our model can infer continuous velocity and standard deviation fields, we can calculate the aleatoric uncertainty on the related flow rate. Additionally, following [13], we use automatic differentiation to obtain pressure gradients and wall shear stress (See Figure 4 and Table 1). The conservative estimate of the aleatoric uncertainty (one standard deviation) is around 20%. Since we only have velocity measurements, obtaining the aleatoric uncertainties related to the other quantities of interest is not viable.

**Table 1:** Average uncertainties (one standard deviation) as percentages of the mean for the volume flow rate near the inlet and outlet (*Q*_*in*_ and *Q*_*out*_), pressure gradient (*∂P/∂y*), and spatially averaged shear stress at the wall 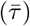 and the shear stress distribution (*τ*).

| | $Q_{in}$ (%) | $Q_{out}$ (%) | $\partial P/\partial y$ (%) | $\bar{\tau}$ (%) | $\tau$ (%) |
| --- | --- | --- | --- | --- | --- |
| Aleatoric | 18 | 20.6 | — | — | — |
| Initialization | 3.4 | 3.1 | 1.6 | 1.1 | 2.9 |
| Model Enhancements | 11.1 | 14 | 8.1 | 6.8 | 11.3 |
| Physical Model | 9.8 | 13.8 | 5.3 | 5.4 | 8.4 |
| Combined | 24.8 | 29.7 | 9.9 | 8.8 | 14.8 |

**Figure 4.**
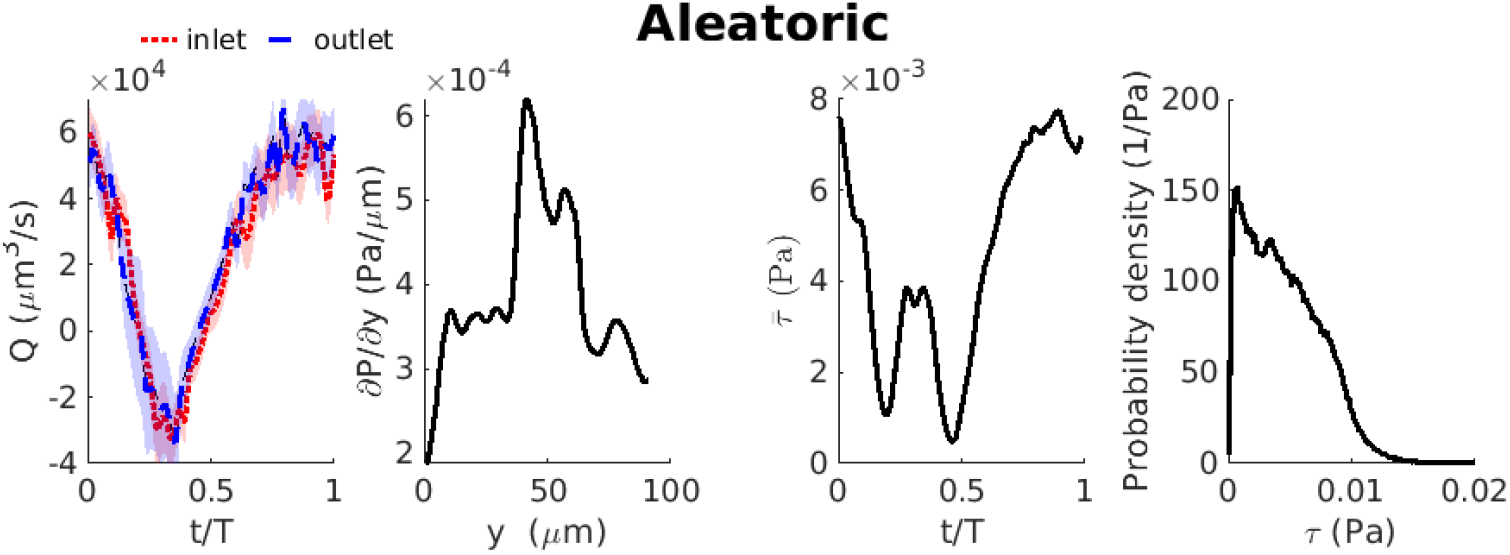
Inferred volume flow rate *Q*, pressure gradient *∂P/∂y*, shear stress at the wall (*τ*) during one cardiac cycle (*T*), and the distribution of wall shear stresses. Since we only have velocity measurements, we can only calculate the aleatoric uncertainty on the flow rate. Shading indicates two standard deviations above and below the mean.

### 4.2. Epistemic Uncertainty

#### 4.2.1. Initialization

We use the deep ensemble [30] method to assess the epistemic uncertainty due to the parameter initialization. Towards this end, we train *M* = 5 models initialized using five different seeds and evaluate their performance in the PTV validation dataset and on the moving boundary conditions.

The mean relative *L*^2^ error over five independent runs on the validation dataset for *u* and *v* are 23.25% and 19.11%, with uncertainty (i.e., one standard deviation) of 0.66% and 0.72%. Similarly, the mean relative *L*^2^ error for the moving boundary conditions for *u* and *w* are 10.75% and 10.45%, respectively, with an uncertainty of 0.75% and 2.05%, respectively (see table A.5).

Additionally, we compute the flow rate, pressure gradient, and wall shear stress for each model and compute the respective mean value and standard deviations using Equation 13 and 14. However, as shown in Figure 5 and Table 1, the uncertainty related to the initialization is much smaller than the other sources of uncertainty (less than 3.5%).

**Figure 5.**
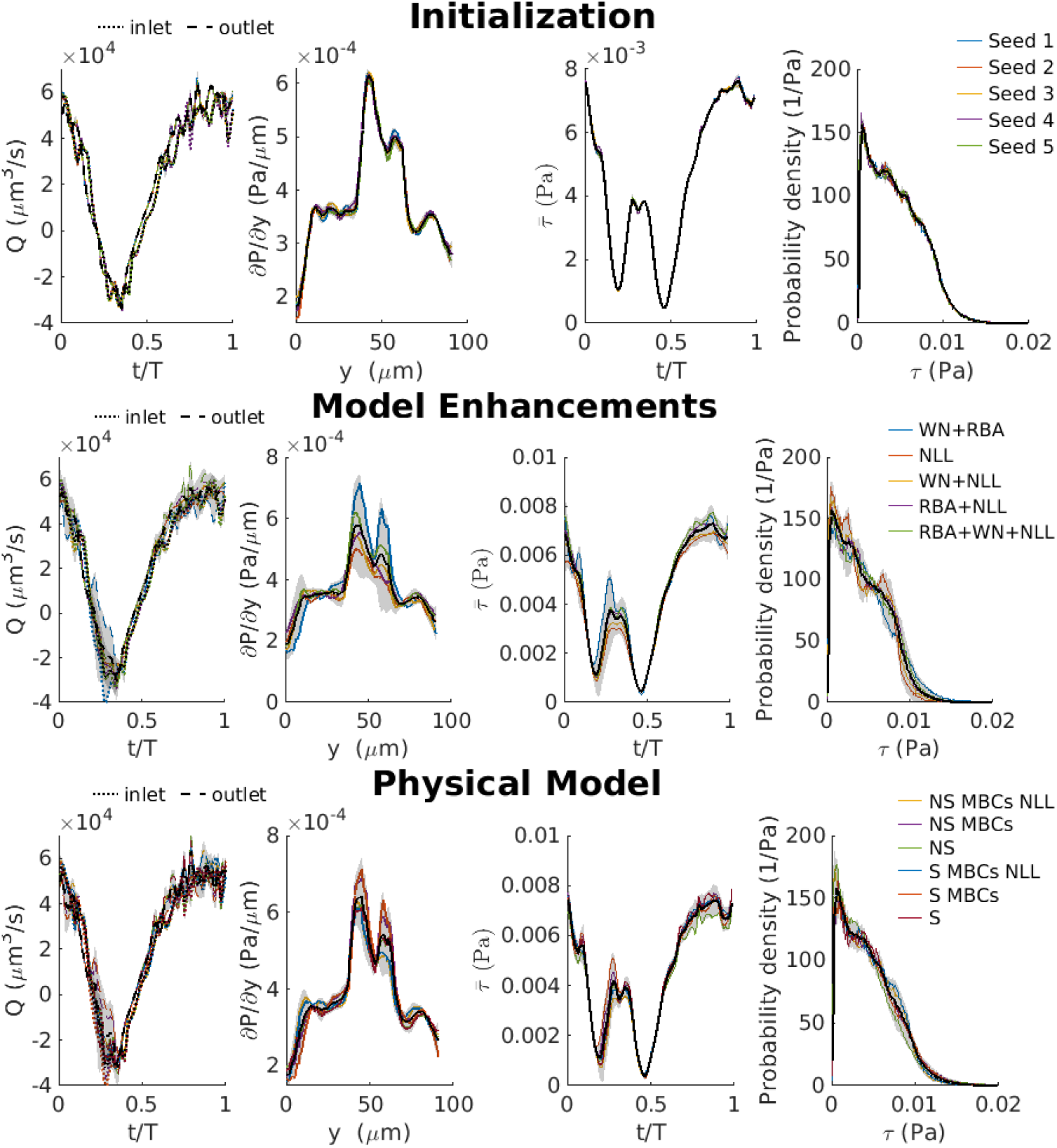
For volume flow rate *Q*, pressure gradient *dP/dy*, and wall shear stress *τ*, the epistemic uncertainty due to the parameter initialization (top row), model enhancement (middle row), and the model uncertainty due to the physical model (bottom row). Shading indicates two standard deviations above and below the mean.

#### 4.2.2. Model Enhancements

Another factor that impacts the inferred solution is the optimization method or model enhancements, introducing possible variations in the predicted outcome. To determine the epistemic uncertainty due to model enhancement, we trained *M* = 5 models using various combinations of weight normalization (WN) [45], residual-based attention (RBA) [16], and Negative Log-Likelihood (NLL).

Table 2 displays the relative *L*^2^ errors on the PTV validation dataset and boundary conditions for the analyzed models. Figure 6 shows that WN is essential for accelerating convergence. Notice that the NLL is crucial for reducing the relative errors in the PTV, particularly for moving boundary conditions, indicating that decomposing the flow field into mean fields and noise is beneficial. Additionally, it can be observed that RBA boosts the model’s performance since it induces residual homogeneity [16].

**Table 2:** Relative *L*^2^ errors as percentages for various model enhancements.

| **Model** | $u_{PTV}$ (%) | $v_{PTV}$ (%) | $u_{BCs}$ (%) | $w_{BCs}$ (%) |
| --- | --- | --- | --- | --- |
| WN+RBA+NLL | 22.54% | 19.14% | 10.89% | 10.80% |
| WN+RBA | 28.97% | 24.78% | 52.20% | 48.10% |
| WN+NLL | 22.95% | 22.50% | 8.35% | 9.61% |
| RBA+NLL | 21.16% | 23.43% | 13.32% | 11.41% |
| NLL | 23.68% | 24.34% | 9.70% | 10.37% |

**Figure 6.**
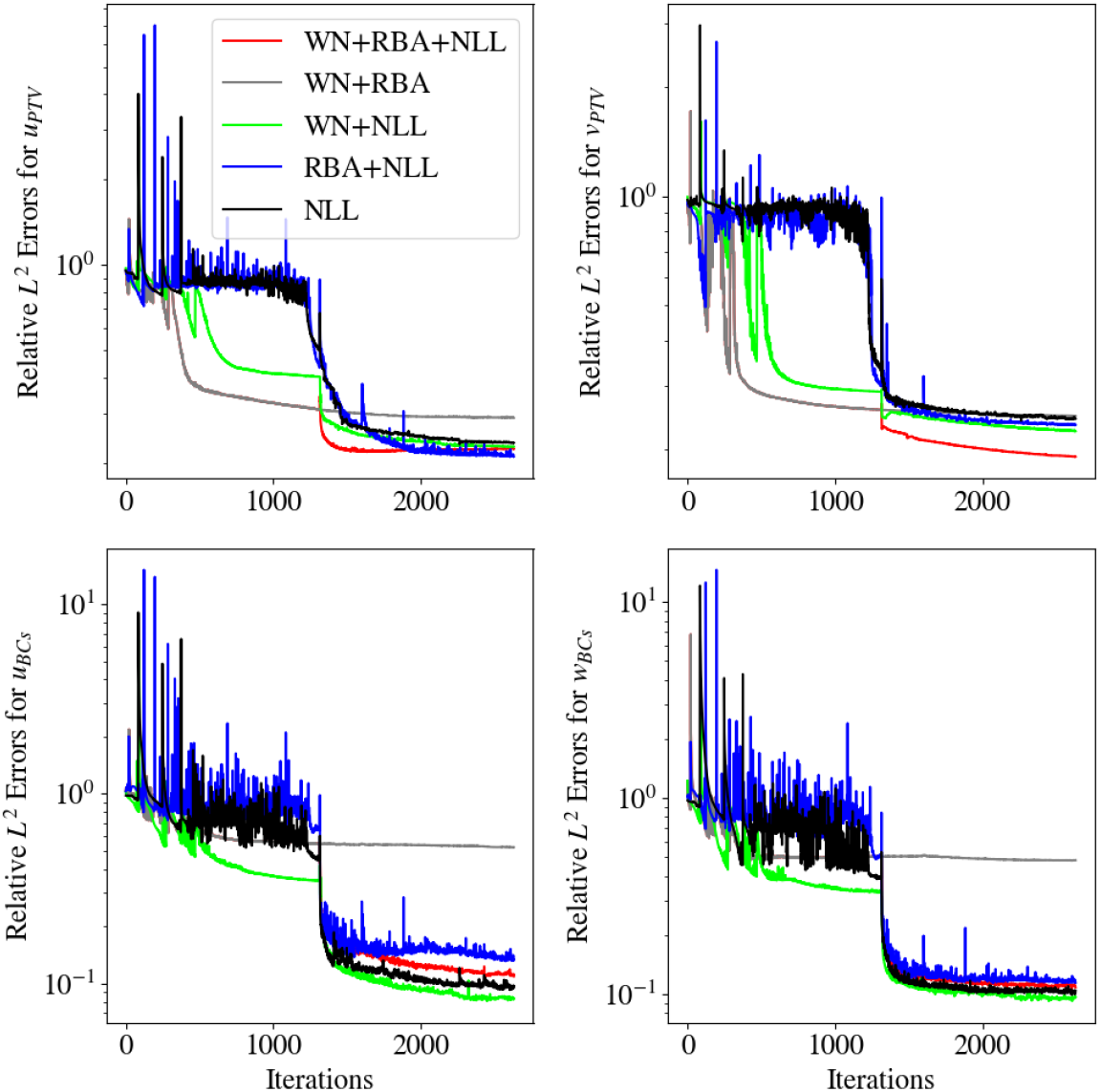
Relative *L*^2^ error convergence history across different model enhancements for the PTV velocity (top row) and moving boundary conditions (bottom row).

Figure 5 (middle row) displays the inferred flow rate, pressure gradient, and wall shear stress for all the analyzed models. The uncertainty (two standard deviations above and below the mean) is represented by the shaded region around the mean field, highlighting the epistemic uncertainty introduced by the model enhancements. The volume flow rate waveform is slightly rougher when RBA is included, which is a result of the model adhering more closely to the noisy experimental training data.

### 4.3. Model Uncertainty

Finally, we analyze the model uncertainty due to the governing equations. To this end, we use the EoM method with *M* = 6 models trained using different physical models (i.e., PDEs, boundary conditions, or other assumptions) that ideally govern the CSF flow. As shown in Table 3, the best-performing models use NLL (i.e., assume that only the mean velocities follow the PDE). These results suggest that the predicted fluctuations (i.e., noise from the experimental data) may be nonphysical. Hence, the proposed decomposition reduces the relative error in the PTV data and boundary conditions.

**Table 3:** Relative *L*^2^ errors as percentages for different physical models and components. N/A stands for not applicable and is related to the models trained using fixed boundary conditions.

| **Model** | $u_{PTV}$ (%) | $v_{PTV}$ (%) | $u_{BCs}$ (%) | $w_{BCs}$ (%) |
| --- | --- | --- | --- | --- |
| NS+MBCs+NLL | 23.14% | 19.52% | 10.73% | 10.41% |
| NS+MBCs | 26.25% | 22.55% | 83.66% | 85.99% |
| NS | 24.58% | 20.84% | N/A | N/A |
| S+MBCs+NLL | 22.54% | 19.14% | 10.89% | 10.80% |
| S+MBCs | 28.97% | 24.78% | 52.20% | 48.10% |
| S | 23.86% | 20.58% | N/A% | N/A |

Additionally, Figures 5 and 7 show that the Stokes and Navier-Stokes models yield comparable performance. This observation aligns with the theoretical assumption that the viscous terms are dominant for *Re*≪1 and the results presented in [13]. However, since the Stokes flow (i.e., equation 3) does not require computing the nonlinear terms, each training iteration is approximately 30% faster than when using NS (i.e., equation 1).

**Figure 7.**
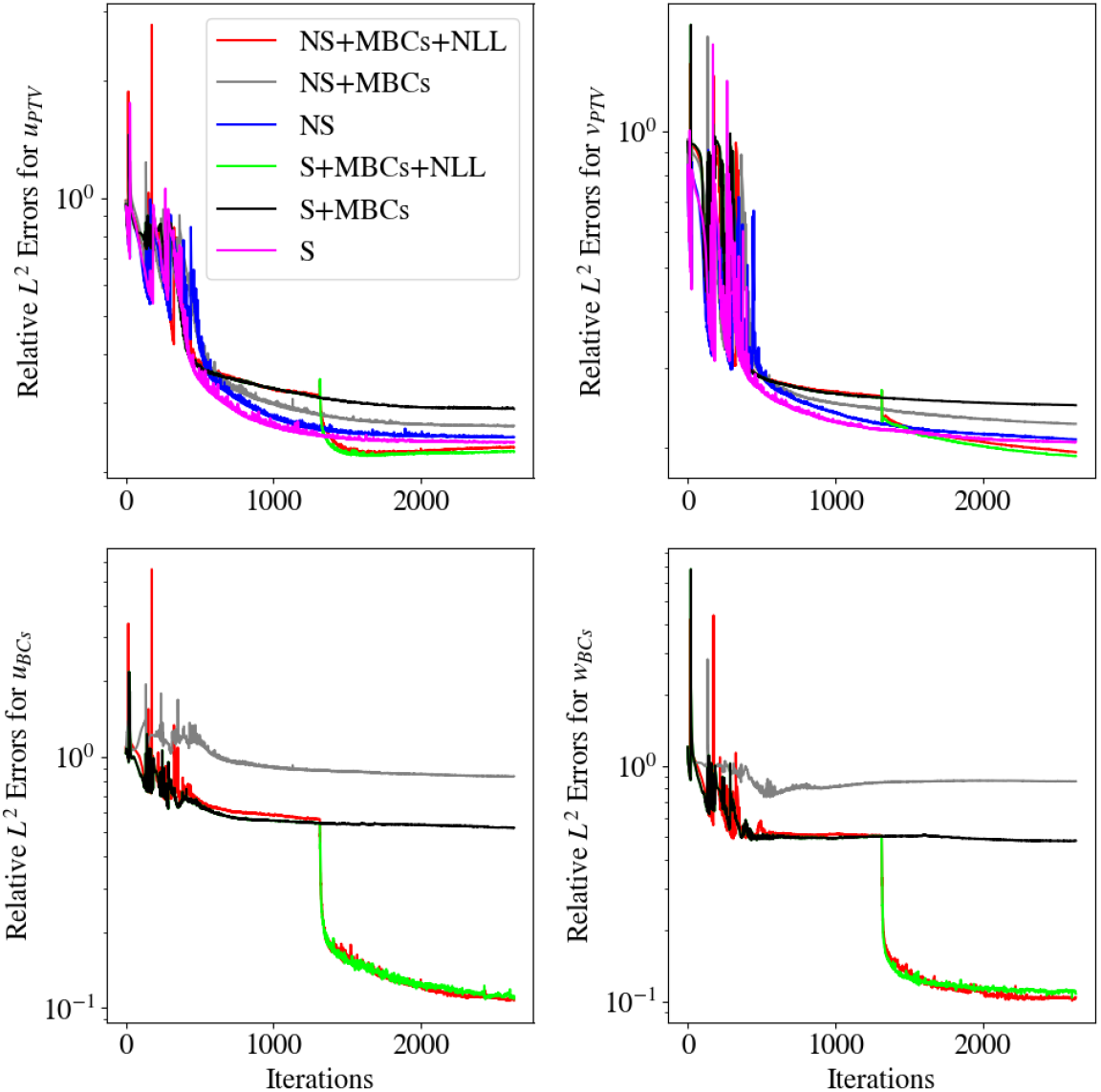
Relative *L*^2^ error convergence history among different physical models for the PTV velocity (top row) and moving boundary conditions (bottom row).

Including moving boundaries (MBCs) did not change the quantities of interest considerably (Fig. 5). The volume flow rate and axial pressure gradient are directly calculated from and related to the downstream velocity *v*, which is aligned with the direction of net flow, so it is unsurprising that the volume flow rate and axial pressure gradient do not change appreciably with the implementation of moving boundaries, which involve motion only in the directions of *u* and *w*. The shear stress magnitude at the wall is calculated from the second invariant of the stress tensor as described in Methods and is thus related to all three flow components. However, as shown in Fig. 5 and Table 1, adding moving boundaries did not change the inferred shear stress substantially either. The cross-stream velocity components, *u* and *w*, are much smaller than the downstream component, *v*. We therefore expect the stress magnitude to be dominated by the component proportional to the wall-normal gradient of the downstream velocity. With respect to the volume flow rate, pressure gradient, and shear stress at the wall, little additional insight is gained by including moving boundaries. For most purposes, using stationary boundaries is appropriate and accurate.

### 4.4. Combined Uncertainty

We combine the model, epistemic, and aleatoric uncertainties as described in equation 15 to get a sense of the overall uncertainty in the model (see Figure 8 and Table 1). The combined uncertainty (one standard deviation) for the volume flow rate is on the order of 30%, with the aleatoric uncertainty accounting for the majority. The pressure gradient, average shear rate, and shear distribution are less uncertain, with combined uncertainties less than 15%, essentially because aleatoric uncertainty cannot be calculated for those quantities, thus it can not be measured. However, the volume flow rate is also more sensitive to each type of epistemic and model uncertainties than the pressure gradient or shear rate. The fact that aleatoric uncertainty is substantially higher than the epistemic and model uncertainties suggests that the most important thing to improve the accuracy of the inferred results is to reduce the uncertainty associated with the experimental measurements. However, when the aleatoric uncertainty is included, the 30% uncertainty is accurate enough for most questions of physiological relevance.

**Figure 8.**
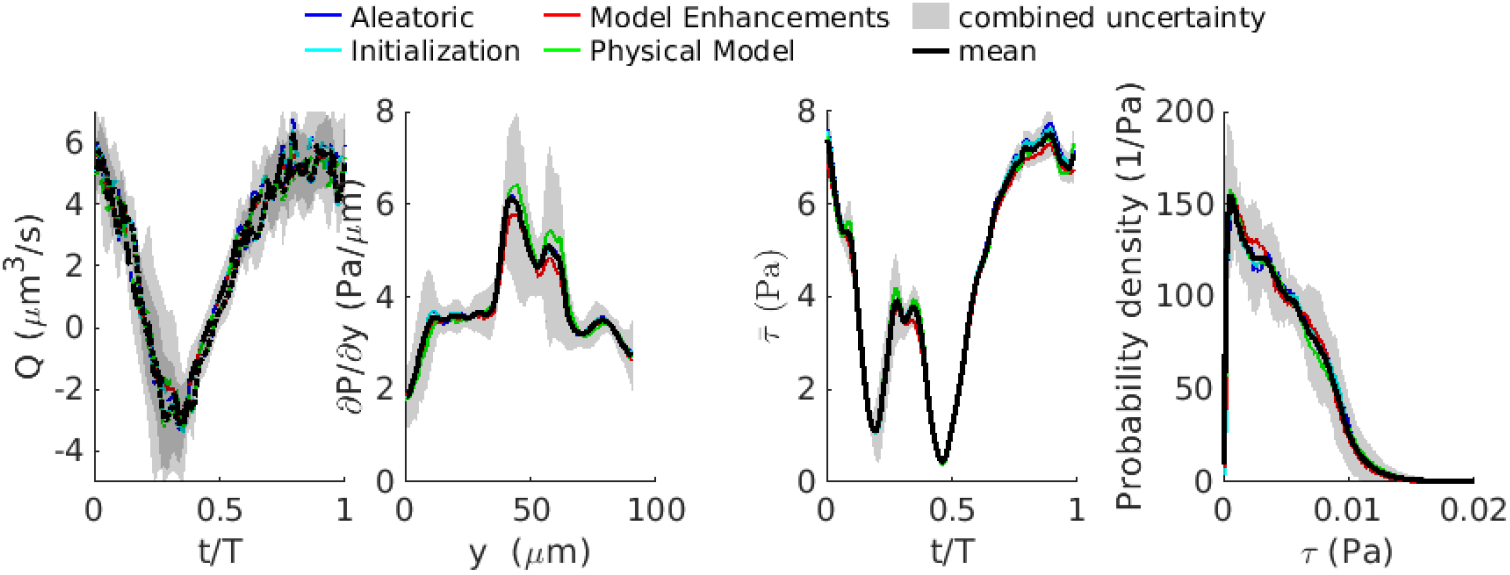
Combined uncertainty in the inferred volumetric flow rate (*Q*), axial pressure gradient (*∂P/∂y*), average shear stress at the wall 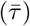, and distribution of shear stresses at the wall (*τ*), for each type of uncertainty (aleatoric, physical model, and epistemic, comprising two uncertainty types: initialization and representation). The average of the inferred quantities of interest from each uncertainty type is shown in black, with shading indicating the combined uncertainty *σ*_combined_ as described in equation 15.

## 5. Discussion and concluding remarks

We demonstrate that using the negative log-likelihood formulation for the loss in artificial intelligence AIV (AIV-NLL) improves the accuracy of AIV inferences and simultaneously quantifies the aleatoric uncertainty in AIV inferred velocity fields and volume flow rates. We also quantify several sources of epistemic uncertainty and model uncertainty in AIV, showing that for volume flow rate, pressure gradient, and wall shear stress, model initialization results in uncertainty less than 3.5%, and modifications to the model (model enhancement) and assumptions regarding the physical model introduce uncertainties less than 15% of the mean. The combined aleatoric, epistemic, and model uncertainty are less than 30% of the mean, demonstrating the accuracy of AIV. We show that using Stokes equations to model the flow reduces the training time by 30%, compared to the full Navier-Stokes equations, with a negligible change in model results.

Boster et al. [13] used an ensemble approach to show that the uncertainty in volume flow rate, pressure gradient, and wall shear stress due to the uncertainty in the 3D domain location (PVS boundaries) was less than 30%, similar to the combined uncertainty we report here for aleatoric, model and epistemic uncertainty, and more significant than any single source of uncertainty alone. This suggests that reducing the uncertainty in the PVS boundaries would have the most significant impact on AIV accuracy and that reducing the uncertainty in PTV measurements would have the second most significant impact. A straightforward approach to reduce the uncertainty in the PVS domain boundaries would be to label the glia limitans (PVS outer boundary) and the endothelial basement membrane (inner boundary), as Raicevic et al. [36] did, rather than relying on the location of dye within the lumen boundaries as Boster et al. [13] did. One simple approach to reduce the uncertainty in the PTV measurements would be to use a smaller field of view. The field of view in the original imaging was 332 by 332 µm, but the AIV analysis was only performed on a smaller subdomain that was 91 by 91 µ. If the original field of view had been zoomed in to this size while maintaining the same number of pixels in the image so that the resolution increased, the uncertainty in the spatial position would have been reduced from 65 nm to 27 nm, and the overall uncertainty would have been reduced from 1.89 to 0.81 µm/s. Of course, two-photon microscopes have resolution limits, but as far as those limits allow, greater magnification substantially reduces uncertainty. Thus, small modifications to the experimental protocol would result in significant improvements in AIV accuracy.

Wall shear stress plays an important role in cardiovascular flows, but little is known about its role in shear in PVSs. Cibelli et al. [46] showed that astrocytes can sense (eliciting a Ca^2^+ response) wall shear stresses that are lower than 0.01 Pa, which is smaller than the highest shear stresses we report here, suggesting that the wall shear stress in physiological glymphatic flows may play an important role in controlling vasomotion and parenchymal perfusion. Cibelli et al. [46] also showed that astrocytes are very sensitive to the magnitude of the wall shear stress. Accordingly, the ability to not only accurately infer wall shear stress but also accurately estimate the uncertainty associated with those inferences becomes extremely important. Cibelli et al. [46] estimated what PVS wall shear stresses would be based on PTV measurements, but shear stress cannot be calculated from PTV directly due to the sparsity of the measurements, and their estimation necessarily involved simplifying assumptions. AIV is the only way to accurately infer 3D wall shear stress measurements from in vivo measurements, and it does so with very little additional computational cost since the derivatives of the flow field are calculated in the network. This work showed that a Stokes flow model with stationary boundaries can be used to infer wall shear stress with very little loss of precision and that the results are insensitive to the initial conditions. These points can inform future efforts to use AIV to infer wall shear stress and further investigate and clarify the role of wall stress in PVSs.

Glymphatic flow has been shown to vary not only with cardiac pulsatility but also at much lower frequencies corresponding to slow vasomotion resulting from neurovascular coupling [5, 28, 47–49]. In order to have sufficient PTV measurements to train the network, we merged velocity measurements from different cardiac cycles together, thus enabling us to infer flow in a single cardiac cycle. Since slow vasomotion cycles are much slower than cardiac cycles, such phase merging may not be required, or at least not over as many cycles. However, since cardiac pulsatility is always present, some approach to disentangle the flow arising from slow vasomotion and that from the cardiac pulsatility would likely be required. Spontaneous slow vasomotion activity is absent in anesthetized mice but present in awake and naturally sleeping mice, which may drive glymphatic flow. However, applying our methods to spontaneous vasomotion would require long imaging sessions of un-anesthetized mice and, therefore, would be logistically challenging. It may be more realistic to infer glymphatic flows due to vasomotion in response to a regular stimulus, as Holstein et al. [5] did. Doing so would provide interesting insight into the relative contributions of cardiac and other vessel wall motion to glymphatic flows.

The AIV-NLL method has several features that distinguish it from the approach used previously in [13]: (1) a modified architecture that predicts the flow fields along with the standard deviations of velocities, (2) neural network re-parameterization using weight normalization, which accelerates convergence, (3) PDE reformulation into Stokes flow, which eliminates the non-linearities and speeds up computation, (4) using negative log likelihood (NLL) instead of mean squared error (MSE) to learn aleatoric uncertainty and further improve model performance, (5) training on uniform batches, which stabilizes the learning process, (6) using fixed global weights based on the scale of the predicted values instead of learning rate annealing [21] to learn the global weights, and (7) using Residual-Based Attention (RBA) as local multipliers, which enables uniform convergence.

These modifications allowed us to reduce the relative *L*^2^ error in *u* by approximately 10% in half the training iterations. Additionally, this approach enabled us to obtain the aleatoric uncertainty, which could not be approximated using the previous approach. Nevertheless, the quantities of interest inferred in this work are similar to those obtained by Boster et al. [13].

## Acknowledgements

All authors acknowledge support from the US NIH National Center for Complementary and Integrative Health (R01AT012312). Additionally, J.D.T., C.W., and G.E.K. acknowledge support from the MURI-AFOSR FA9550-20-1-0358 project, the ONR Vannevar Bush Faculty Fellowship (N00014-22-1-2795) and the DOE SEA-CROGS project (DE-SC0023191). K.A.S.B and D.H.K. acknowledge the support from and from the BRAIN Initiative of the US National Institutes of Health (U19NS128613) and from the US Army (MURI W911NF1910280). Finally, we thank Zongren Zou, from Brown University, for his insights on uncertainty quantification.

## Appendix A. Implementation Details

### Appendix A.1. Training Domain

**Figure A.9:**
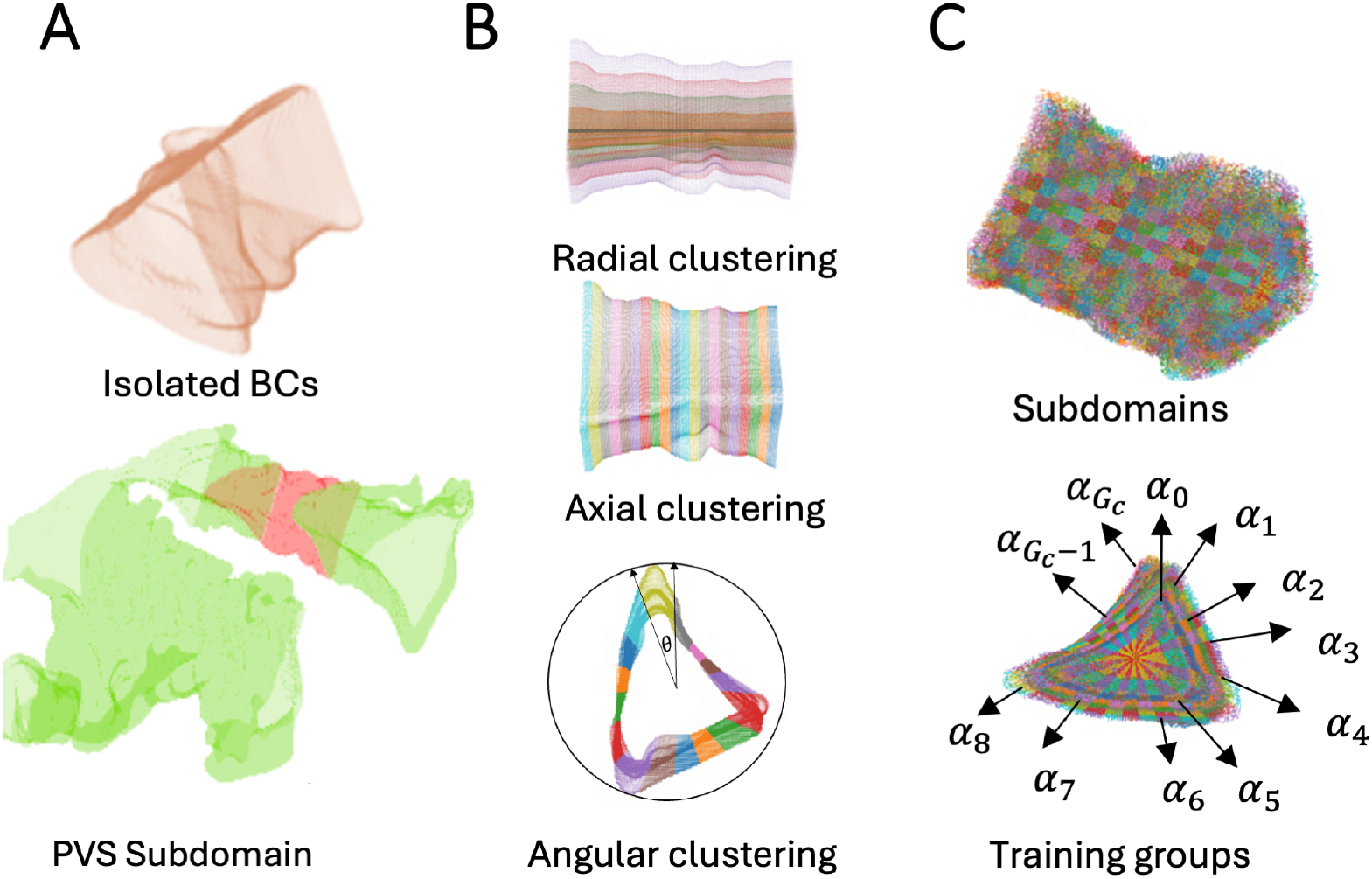
(A) We obtain our boundary conditions (BCs) by isolating the PVS subdomain. (B) We cluster the BCs based on their spatial and temporal coordinates and obtain *N*_*θ*_ angular, *N*_*y*_ axial subgroups. After that, we generate points inside the domain by shrinking and combining boundary conditions. We classify these points into *N*_*r*_ radial sections. Finally, we tile them and classify them into *N*_*t*_ temporal subgroups. (C) Based on the classification, we obtain information on subdomains that define training groups used to generate ordered mini-batches.

Following [13, 16], we train our model using 30% of the available PTV data and use the remaining unseen data for validation. Given the scarce amount of experimental data, we use full-batch training at every iteration. However, we use ordered mini-batches by clustering the BCs and collocation points. As described in Section 3.1.2, we obtain boundary conditions by isolating a subdomain of PVS and defining the boundary motion. Since the boundary points delimit the training domain, we follow [16] and generate our collocation points (to evaluate PDEs) from the BCs; see Figure A.9(B). First, we split our boundaries into 25 angular subregions (*N*_*θ*_) and 15 axial subregions (*N*_*y*_). Then, we shrink and combine these subregions to get an equally spaced domain of 200 layers. After that, we split our domain into seven radial groups (*N*_*r*_), tile them 101 times, and split the time domain into 20 temporal subregions (*N*_*t*_). Consequently, the total number of training groups (*G*_*c*_) for the Navier-Stokes (NS) equations is

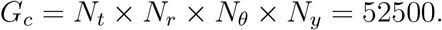

Similarly, we split the boundary conditions with *N*_*θ*_ = 25, *N*_*y*_ = 15, and *N*_*t*_ = 15, so the number of training groups is

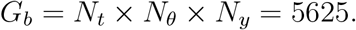

Notice that the total number of collocation points *G*_*c*_ × *N*_*b*_ = 4.98 × 10^6^ and boundary condition points *G*_*b*_ ×*N*_*b*_ = 5.34 × 10^5^ approximately matches the number of training points used in the previous study (i.e., 5 × 10^6^ and 5 × 10^5^) [13].

Following this method, we obtained 95 ordered mini-batches used to train our model. As described in [16], by this approach, each mini-batch contains information about the whole domain, which accelerates and stabilizes the model convergence.

### Appendix A.2. AIV-NLL

The training data comes from experimental measurements and thus has inherent uncertainty. A conservative estimate of the uncertainty is 1.89 µm/s [13]. Though Boster et al. showed the uncertainty in the quantities of interest due to uncertainty in the boundary location, they used the same velocity measurements for training, and the impact of the uncertainty in the velocity measurements was not investigated. Here, we use AIV-NLL to infer the uncertainty in the velocity measurements and how that uncertainty affects the calculated volumetric flow rate.

Given the different scales of the predicted values, the mean fields 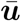 are approximated as follows:

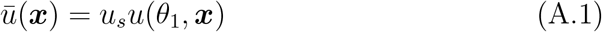

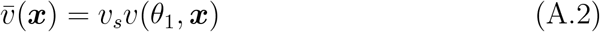

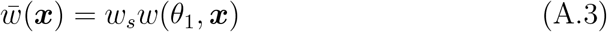

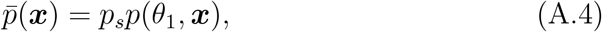

where *u, v, w, p* are the outputs of the first neural network, which approximates the mean velocity and pressure fields, scaled by *u*_*s*_, *v*_*s*_, *w*_*w*_ and *p*_*s*_, respectively. The corresponding standard deviations *σ*_*u*_, *σ*_*v*_ and *σ*_*z*_ for the velocity in *x, y*, and *z* are computed as

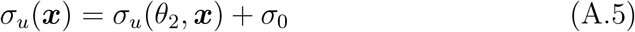

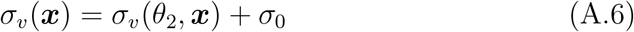

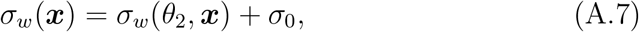

where *σ*_*u*_(*θ*_2_, ***x***), *σ*_*z*_(*θ*_2_, ***x***) and *σ*_*w*_(*θ*_2_, ***x***) are the predicted standard deviations and *σ*_0_ is a positive number that defines a lower bound.

We evaluate our model performance using the relative *L*^2^, which is defined as:

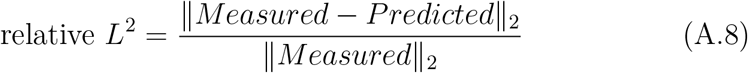

### Appendix A.3. Additional Model Enhancements

#### Appendix A.3.1. Weight Normalization

Weight Normalization is a re-parameterization technique that accelerates convergence in PINNs [12]. In this scheme, the weight vectors’ length and direction are decoupled to be trained separately [45]. Each neuron output is defined as follows:

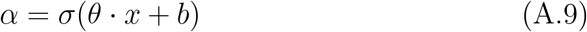

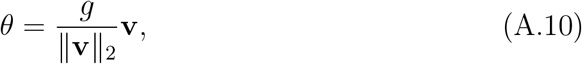

where *α* is the neuron output, *σ* is the activation function, *x* is the input vector, *θ* is a weight vector, and *b* is the bias. As shown in equation A.10, the weight vector *θ* is redefined in terms of new trainable parameters, **v** (direction) and *g* (length). Notice that ||*θ*|| = *g*, so this re-parameterization allows us to decouple the weight’s length and direction, which speeds up the model convergence. Since *g* is a scalar, this modification induces minimal computational overhead [45].

#### Appendix A.3.2. Residual Based Attention (RBA)

Training neural networks often involves the challenge of residuals (i.e., point-wise errors) being overlooked when calculating the cumulative loss function, which typically involves the summation or mean of the residuals [16, 23]. To address this issue, several studies have proposed scaling the loss terms using local multipliers [16, 20]. These local multipliers, such as residual-based attention (RBA) weights [16] and self-adaptive weights [20], have demonstrated remarkable performance in physics-informed neural networks (PINNs) and other supervised learning tasks. By balancing the contribution of specific training points within each loss term, these weights induce residual homogeneity [17].

RBA weights are based on the exponentially weighted moving average of the residuals. Since the loss residuals contain information about high error regions, the resulting multipliers act as an attention mask, helping the optimizer focus on capturing the spatial or temporal characteristics of the specific problem [16].

The update rule for RBA for any training point *i* on iteration *k* is given by

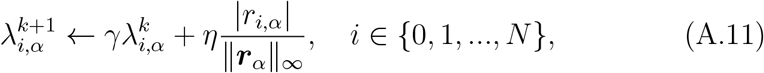

where *N* is the number of training points, *r*_*i,α*_ is the residual of the loss term (*α*) for point *i*, and *η* is a learning rate and *γ* is a memory rate that enables the model to eventually “forget” the previous iterations. This convergent linear homogeneous recurrence relation bounds our RBA from zero to one (*λ*_*i,α*_ ∈ [0, *η/*(1 − *γ*)]) [16, 17, 35].

### Appendix A.4. Training

We approximate the mean 3D velocities (*u, v, w*) and pressure (*p*) as described in equations A.1, A.2, A.3 and A.4, respectively. To ensure that the model outputs have the same order of magnitude, we scale them using (*u*_*s*_, *v*_*s*_, *w*_*s*_, *p*_*s*_) = (0.1, 1, 0.1, 100). Similarly, for the predicted uncertainties, we use a constant *σ*_0_ as a lower bound. *σ*_0_ = 1.9*µm/s* is chosen based on the minimum uncertainty related to the measurement technique.

We train our model during 2632 epochs (approximately 2.5*e*5 iterations) using Adam Optimizer. During the first half of training, we focus on learning the mean fields using *MSE* in the data and boundary loss (i.e., equations 9, 10). In this stage, we use an auxiliary loss to initialize the uncertainties to *σ*_*u*_ = *σ*_*v*_ = *σ*_*w*_ = 1. Notice that when ***σ*** = **1** the mean NLL reduces to *MSE*.

In the second stage, we learn the corresponding standard deviations by using NLL in the data and boundary loss. Notice that in the NLL formulation (i.e., equation 8), we use a constant *C* = log(*σ*_0_) that ensures that the loss criterion for any point is always positive.

During training, we used global weights *m*_*α*_ to balance the contribution of each loss component. The specific global weights are chosen so that the relative contribution of all loss terms is in the same order of magnitude. The specific values were selected following the previous study [16] and are described in Table A.4.

**Table A.4:**
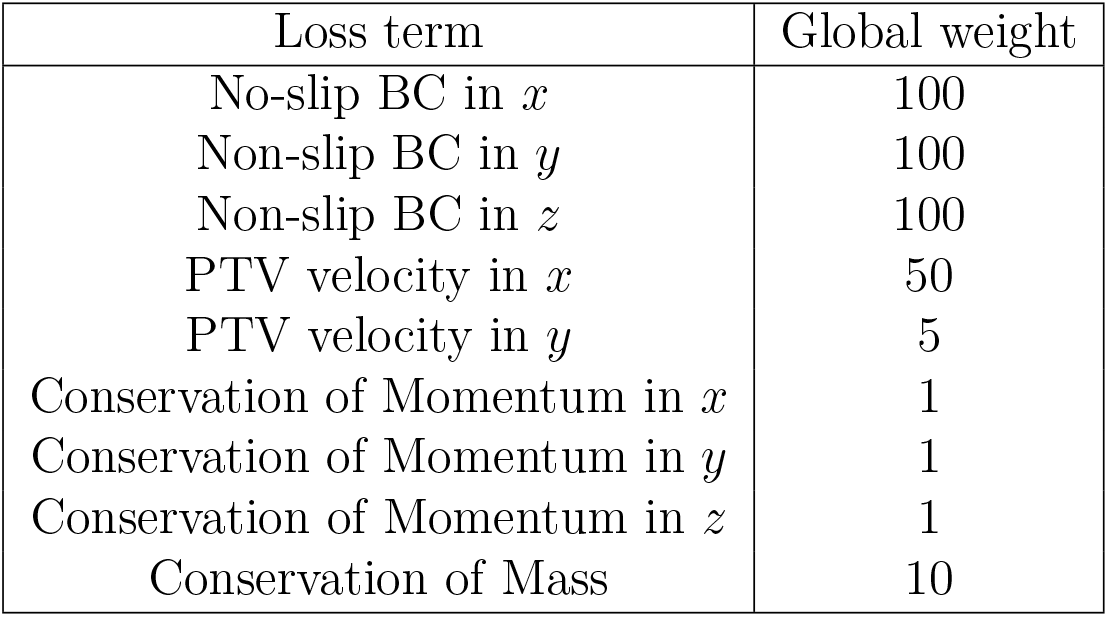
Global weights for AIV-NLL.

| Loss term | Global weight |
| --- | --- |
| No-slip BC in $x$ | 100 |
| Non-slip BC in $y$ | 100 |
| Non-slip BC in $z$ | 100 |
| PTV velocity in $x$ | 50 |
| PTV velocity in $y$ | 5 |
| Conservation of Momentum in $x$ | 1 |
| Conservation of Momentum in $y$ | 1 |
| Conservation of Momentum in $z$ | 1 |
| Conservation of Mass | 10 |

### Appendix A.4.1. Hyperparameter Settings

The remaining implementation details are selected based on our previous studies [13, 16]. In particular, for both neural networks, we use a multilayer perceptron (MLP) with eight hidden layers and 200 neurons per layer using the *sin*(·) activation function. The initial learning rate is set to 2 ×10^−3^, which decays to a final learning rate of 1.5×10^−4^ with a decay rate of 0.9. We use RBA weights to scale the point-wise contribution inside the loss term using a decay rate *γ* = 0.999 and learning rates of 0.2 for PTV and Navier-Stokes and 0.02 for BCs. As described in [16] we use a reduced learning rate for the boundaries due to their high uncertainty as reported in [13]. The machine learning framework used is Jax, and the computations are performed on an Nvidia A100 GPU.

### Appendix A.4.2. Additional Results

**Table A.5:** Relative L^2^ errors as percentages for various parameter initialization. The mean and standard deviation for each component: *u*_*P T V*_ mean = 23.25%, std = 0.66%; *v*_*P T V*_ mean = 19.11%, std = 0.72%; *u*_*BCs*_ mean = 10.75%, std = 0.25%; *w*_*BCs*_ mean = 10.45%, std = 0.65%.

| Seed | $u_{PTV}$ (%) | $v_{PTV}$ (%) | $u_{BCs}$ (%) | $w_{BCs}$ (%) |
| --- | --- | --- | --- | --- |
| 1 | 23.95% | 18.63% | 10.39% | 10.01% |
| 2 | 22.54% | 19.14% | 10.89% | 10.80% |
| 3 | 22.57% | 20.10% | 11.06% | 9.97% |
| 4 | 24.10% | 18.07% | 10.15% | 11.58% |
| 5 | 23.09% | 19.63% | 10.90% | 10.09% |

**Figure A.10:**
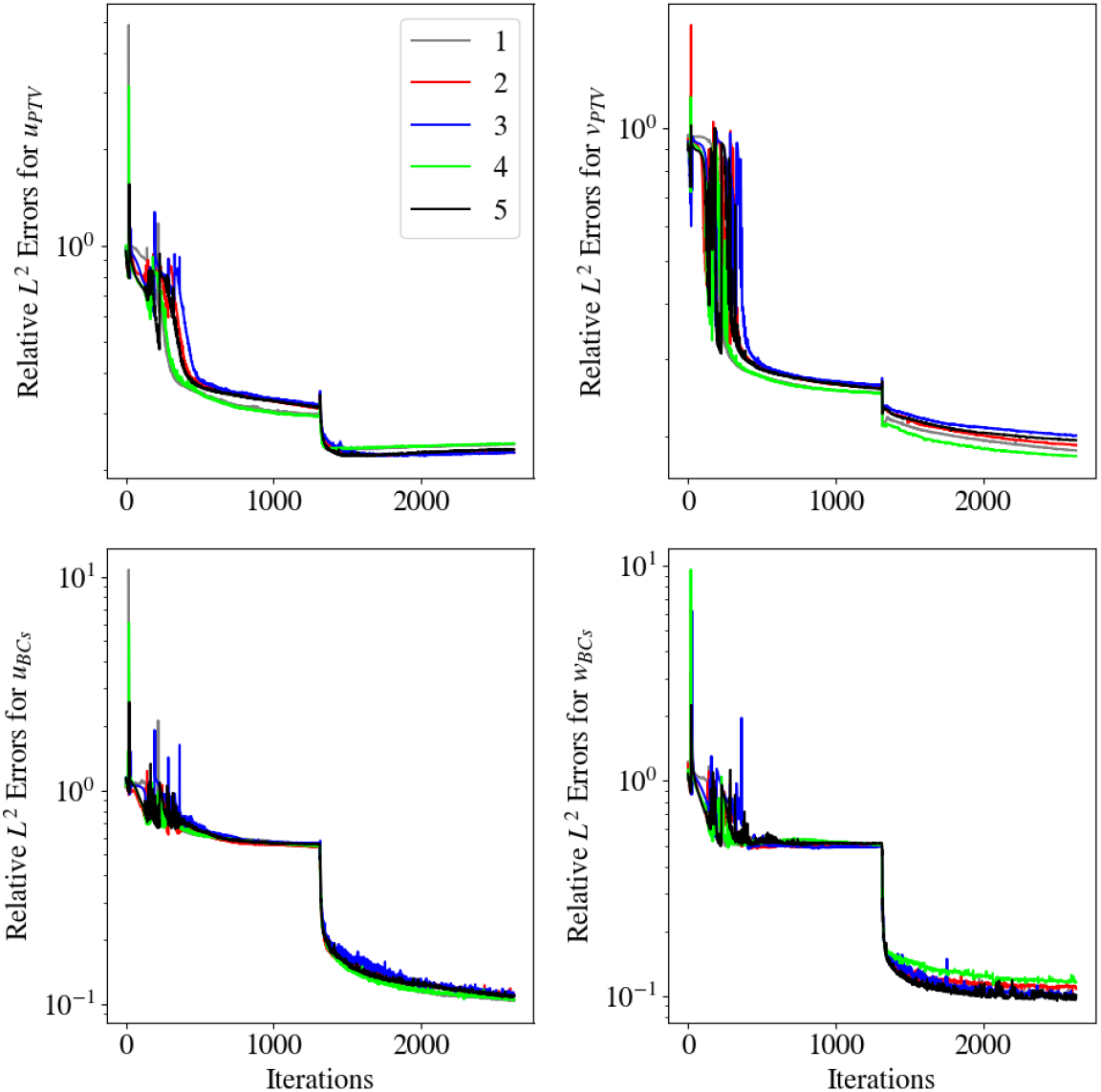
Relative *L*^2^ error comparison among different parameter initialization for the PTV velocity (top row) and moving boundary conditions (bottom row)

